# G-quadruplexes catalyze protein folding by reshaping the energetic landscape

**DOI:** 10.1101/2024.12.18.629195

**Authors:** Zijue Huang, Kingshuk Ghosh, Frederick Stull, Scott Horowitz

## Abstract

Many proteins have slow folding times *in vitro* that are physiologically untenable. To combat this challenge, ATP-dependent chaperonins are thought to possess the unique ability to catalyze protein folding. Performing quantitative model selection using protein folding and unfolding data, we here show that short nucleic acids containing G-quadruplex (G4) structure can also catalyze protein folding. Performing the experiments as a function of temperature demonstrates that the G4 reshapes the underlying driving forces of protein folding. As short nucleic acids can catalyze protein folding without the input of ATP, the ability of the cell to fold proteins is far higher than previously anticipated.

**Significance Statement:** How folding of proteins occurs en masse in the cell is still a daunting unsolved problem. Many proteins have complicated and difficult folding trajectories, with *in vitro* folding times that are physiologically untenable. The acceleration of protein folding to physiologically relevant timescales is a biologically essential function thought to be accomplished by a small set of ATP-dependent chaperonins. In this work, we surprisingly show that small nucleic acid sequences containing G-quadruplexes can catalyze protein folding and reshape protein folding energy landscapes. As a result, the capacity for accelerating protein folding in the cells is far higher than previously suggested, potentially explaining the accommodation of large number of proteins with physiologically unreasonable folding times.

## Introduction

Molecular chaperones are molecules that maintain a healthy protein folding state of the cell, termed proteostasis. Chaperones are often broken into two categories, ATP-independent chaperones (often termed holdases), and ATP-dependent chaperones (often termed foldases). ATP-independent chaperones such as small heat shock proteins are typically thought to bind to aggregation-prone sections of protein and prevent their further aggregation, but not directly aid folding. Instead, they later transfer their clients to ATP-dependent chaperones for refolding. ATP-dependent chaperones, such as Hsp60, Hsp70, and Hsp90, hydrolyze ATP to drive large conformational changes that can directly aid in unfolding and refolding proteins (1). While the exact mechanisms that ATP-dependent chaperones use to aid protein folding are in many cases still under debate, GroEL in particular has shown the ability to accelerate protein folding for proteins encapsulated within its cavity, and this folding acceleration has been hypothesized to be a unique feature of chaperonin cavities (2). While other chaperones allow folding while bound to their surface, so far folding-while-bound has been found to slow down protein folding (3). To our knowledge, true protein folding catalysis with increased rates in both forward and reverse directions has so far only been demonstrated by the GroEL cage.

G-quadruplexes (G4s) are four-stranded nucleic acid structures that can form with all four strands in the same direction (parallel), in opposing directions (anti-parallel) or with three in the same direction and one in the opposite direction (3+1 hybrid) (Fig 1A). In eukaryotes, RNA G4s preferentially fold under stress conditions and unfold when the stress dissipates (4). Recently, we and others have shown that G4s are powerful modulators of proteostasis (5–7). Unlike ssRNA and dsDNA that only helped protein folding indirectly, we found that RNA and DNA G4s can rescue protein folding intermediates and thus help a protein achieve its native state. In cells where degradation and translation had been paused, the primary effect of added RNA G4s on the test protein was to improve its folding (5). However, the mechanism of how G4s affect protein folding remained unresolved.

**Figure 1.**
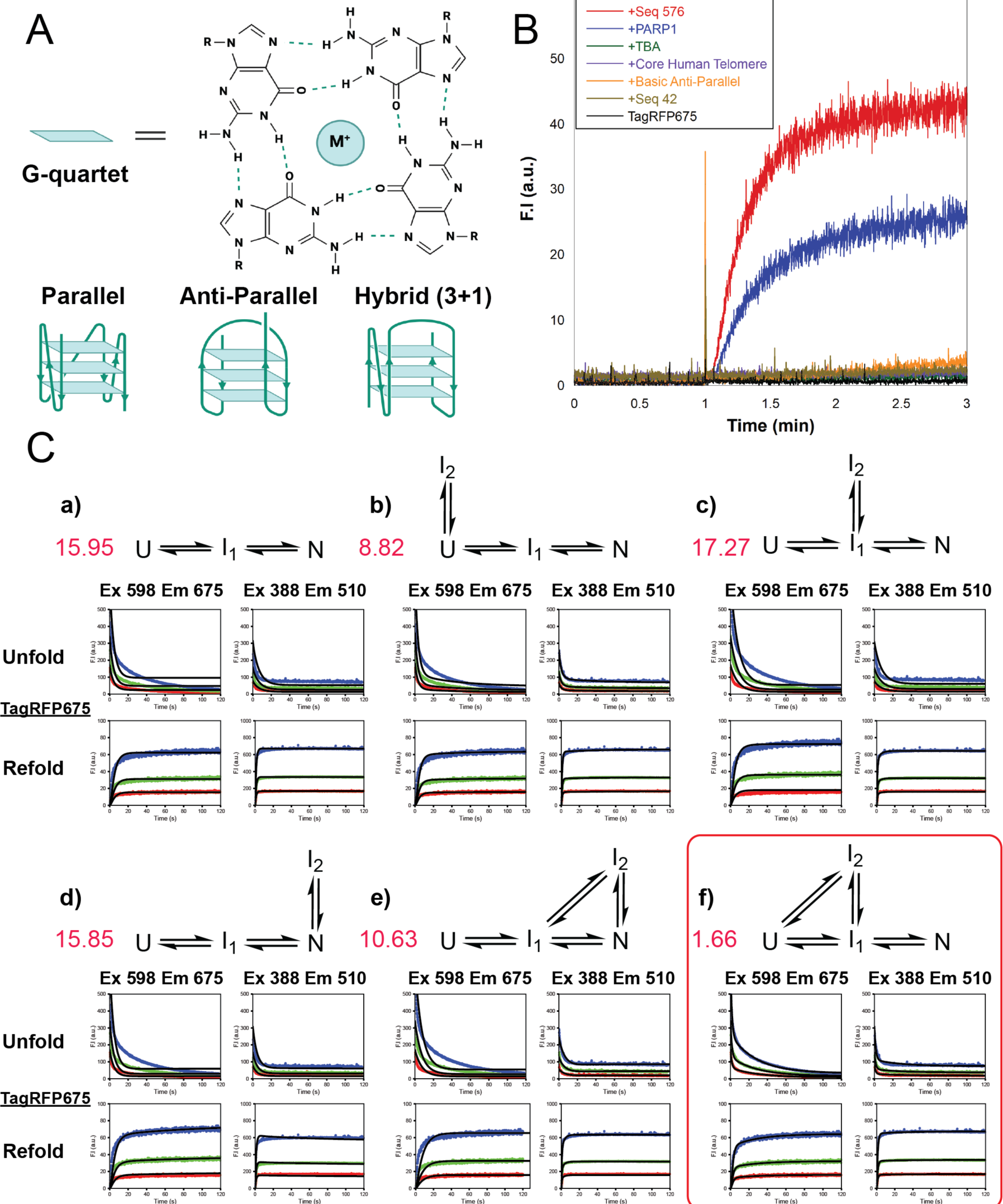
G4 structure and TagRFP675 folding on its own. A. G4 structure. A G-quartet contains four guanines on the same plane interacting via Hoogsteen hydrogen bonding, stabilized by a cation. The arrow direction on the unimolecular G4 shows different conformations. B. Fluorescence recovery of TagRFP675 depends on the different G4 topologies. Seq 576 is an parallel G4, PARP1 is 3+1 hybrid G4. TBA, core human telomere, and basic anti-parallel are all anti-parallel G4s. Seq42 and the absence of DNA are served as the negative controls. The ssDNA:TagRFP675 ratio for all is 2:1. C. Step-wise global fitting of TagRFP675 folding mechanism (a-f) at 20 °C using Kintek Explorer. f is the best fit mechanisms (*X*^2^/dof = 1.66). TagRFP675 concentrations are 0.5 (red), 1 (green), and 2 (blue) μM for unfolding, and 0.25 (red), 0.5 (green), and 1 (blue) μM for refolding.

In this work, we performed folding and unfolding kinetics experiments in the presence and absence of a G4. Surprisingly, we discovered that the G4 acts as a protein folding catalyst, increasing both the rate of folding and unfolding of the protein to both an on-pathway intermediate and its native state. The addition of G4 also effectively removes accumulation of off-pathway intermediates. We performed these experiments as a function of temperature and determined that the G4 affects the underlying thermodynamic driving forces of protein folding differently for different steps of the folding pathway. That a short nucleic acid sequence without a chaperonin cavity can act as a folding catalyst suggests that nucleic acids could play a much larger role in actively and globally modulating protein folding times in the cell than previously considered.

## Results

### Topology dependence of G4 foldase activity

Before determining the detailed mechanism of G4s in protein folding, we checked whether the folding capacity of G4s is dependent on G4 topology type. Therefore, we performed protein folding experiments with TagRFP675 in the presence of either parallel, anti-parallel, or mixed 3+1 G4s. Seq 576 (parallel) enhanced folding the most, follow by PARP1 (3+1 hybrid). Anti-parallel sequences showed no chaperone activity. This result closely matches our previously observed results for the topology dependence of G4s in preventing protein aggregation (6, 8). We chose to continue with Seq576, as it reproducibly forms a parallel G4 structure with a high degree of thermal stability and remains a parallel G4 over the entire temperature range examined here (6).

### Mechanism of the folding of TagRFP675

Before investigating the role of the G4 in protein folding, the protein itself needed its folding mechanism elucidated. TagRFP675 has previously been shown to fold with at least one fluorescent intermediate (5, 9). Therefore, we started by fitting the simplest three-state model, an unfolded protein (U) that has no fluorescence signal, an on-path fluorescent intermediate (I_1_), and native protein (N) with the respective excitation emission wavelengths of 388/510 nm and 598/675 nm. The temperature was set at 20 °C. Of note, the U step contains any states prior to the fluorescent intermediate, and therefore could potentially also include any earlier non-fluorescent intermediates. The folding of N was fitted as a double exponential due to a burst at the beginning followed by a plateau, suggesting the presence of an intermediate. The folding of I_1_ was fitted as single exponential. As can be seen in Fig. 1B, a), this simple three step mechanism poorly describes data (χ^2^/dof = 15.95). We then systematically increased the complexity of the model by including an off-pathway (I_2_) intermediate to represent trapped intermediates (10) or aggregates (Fig. 1B, b-f). A good agreement with data was achieved (χ^2^/dof =1.66) in which the off-pathway I_2_ interconverts with both U and I_1_, but not N. More aggregation steps were also attempted but goodness of fit did not improve by adding further complexity (Data not shown).

In addition to the mechanism, we could determine rate constants for these steps. At 20 °C, the rate of U to off-pathway I_2_ (0.73 s^-1^) is approximately 8-fold higher compared to on-path folding from U to I_1_ (0.09 s^-1^). The return from I_2_ to I_1_ and I_1_ to U is very slow (both <0.1 s^-1^), suggesting that much of the protein was trapped off-pathway. Interconversion between on-pathway I_1_ and N was faster and of similar rates in each direction (0.21 s^-1^ forward and 0.44 s^-1^ reverse). These rates yield an effective folding time of approximately 17 seconds in reasonable agreement with theoretical estimates (18 to 38 seconds, see methods for more) using predictive models of protein folding kinetics (11, 12). With the protein folding mechanism characterized, we could now interrogate the effects of adding a G4.

### Mechanism of G4 assisted folding

As a control, we first tested whether addition of G4 to the experiment required a change in the modelling to account for its presence. We therefore performed global fitting of the TagRFP675 mechanism by itself (Fig. 1B, f) but using the folding and refolding data in the presence of the G4. As DNA Seq 576 has previously been showed to enhance TagRFP675 folding in *E. coli*, here we use this G4 in all our kinetic analysis (Fig. 1A). Using data collected in the presence of G4 but without inclusion in the model, the data was poorly fit (χ^2^/dof = 7.74), showing that the effects of the G4 must be accounted for (Fig. S1).

To determine how the G4 affects the folding of TagRFP675, we began modeling the mechanism of folding including folding in the presence of G4. This mechanism includes steps accounting for the unfolded-G4 complex (UG4), on-pathway intermediate bound to G4 (I_1_G4) and native state bound to G4 (NG4). We systematically tested each model, increasing complexity one step at a time to create the least complicated model that could account for the data (Fig. 2A, Fig. S2). The model that fits our G4-assisted kinetics invokes a folding-while bound model (Fig. 2B). The protein proceeds through folding while continually bound to the G4.

**Figure 2.**
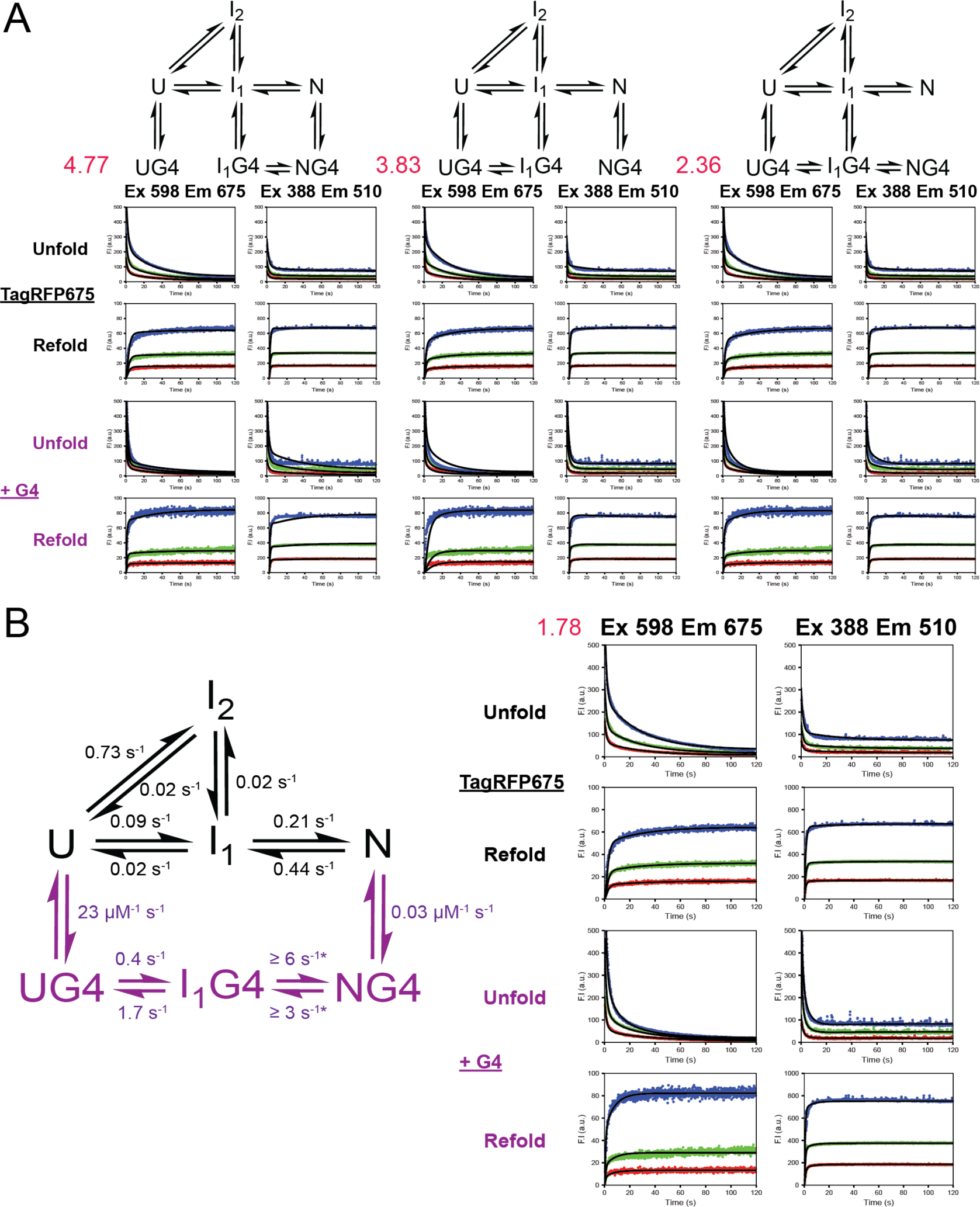
Global fitting of G4-catalyzed mechanism at 20 °C. Purple denotes the mechanism and data collected in the presence of G4, black denotes the mechanism without G4 and data collected in the absence of G4. A. Folding while bound mechanism step-wise fitting. B. Best fit TagRFP675 with G4 folding mechanism (*X*^2^/dof = 1.78) and fit fluorescence kinetic assays. Off-pathway step from I_1_ is required for a good fit even though the reaction rate is very small. Steps that are not labeled on the scheme are not well defined by the data. In case when only lower limits can be defined (asterisk), good fit can be achieved as long as the ratio of forward and reverse is maintained. All rate constants obtained from global fitting can be found in Table S2. All are 2:1 ratio of G4:TagRFP675, TagRFP675 concentrations are 0.5 (red), 1 (green), and 2 (blue) μM for unfolding, and 0.25 (red), 0.5 (green), and 1 (blue) μM for refolding.

We noted that no off-pathway I_2_G4 state was required for fitting, which prompted us to assess whether I_2_ was required in the fitting at all when G4 was present. Fitting only the data in the presence of G4 showed that when G4 was present, the off-pathway I_2_ state was no longer needed to achieve a good fit (Fig. S3). This result suggests that one property of the G4-assisted folding is to effectively prevent the formation of off-pathway intermediates and aggregates.

Next, we turned to folding and unfolding rate analysis. Using all of our fit data, we constrained all rate constants by 95% confidence contour analysis (Fig. S4-S6). The binding of U to the G4 is relatively fast compared to the other available paths, which effectively shunts the protein to fold while bound to the G4 and effectively removes protein from forming the off-pathway I_2_ state, explaining why I_2_ was no longer required for fitting. Surprisingly, the rates of both folding and unfolding of each step increased in the presence of the G4 by at least 5-fold. As the rate increases were roughly proportional in both directions, these results indicate that the G4 catalyzes protein folding.

### Temperature-dependence of Folding

To determine the underlying energetic driving forces of the protein folding catalysis by the G4, we performed these same folding experiments with and without G4 at five more temperatures (Tables S1, S2, and Fig S6). Then, we used the Eyring equation to determine changes in enthalpy (ϕλH^‡^), entropy (ϕλS^‡^), free energy (ϕλG^‡^), and potentially heat capacity (ϕλCp^‡^) for each transition state (Fig 3A, Table 1, Figs S7, S8). In all but one step, the data was well-explained without changes in heat capacity. The U to I_1_ transition, however, showed distinct curvature in the Eyring plot, necessitating the inclusion of heat capacity for that step. Two primary trends emerge from the thermodynamic parameters. For all transitions, the addition of G4 caused an increase in the enthalpic barrier, but the G4 also caused a decrease in the entropy barrier. In all cases other than U to I_1_, the entropy barrier decrease is larger in magnitude than the enthalpy barrier increase. For U to I_1_, transition is not primarily the entropy barrier decrease, but a change in heat capacity.

**Figure 3.**
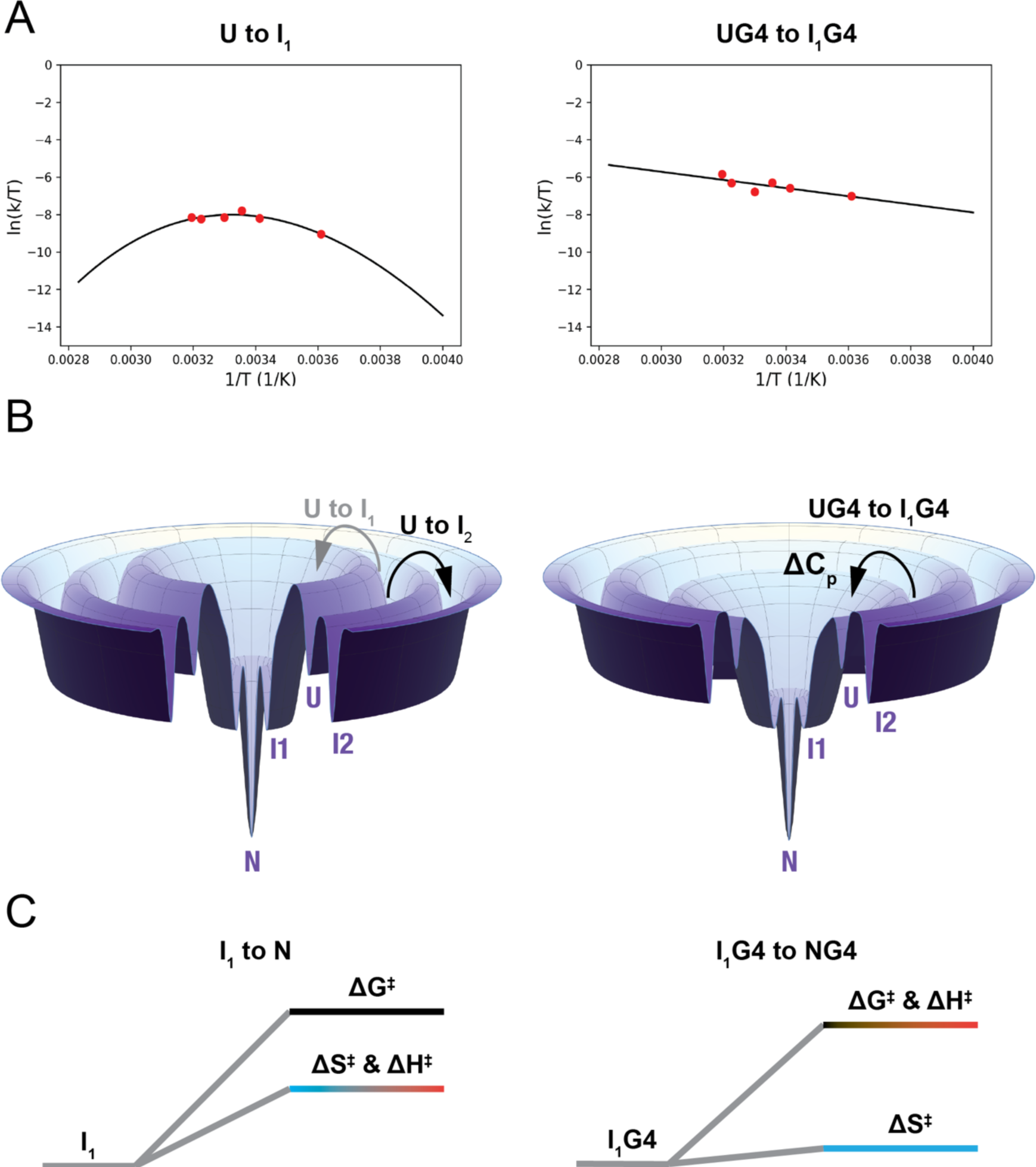
Temperature dependence of. A. TagRFP675 Eyring plot example in the absence (left) TagRFP675 and in the presence of G4 (right). B. TagRFP675 folding energy landscape in the absence (left) and in the presence of G4 (right) at 20 °C. From U to I_1_, folding is dominated off-pathway protein accumulation without G4. Folding while bound to the G4 is faster, outcompeting the off-pathway state due to heat capacity change. C. Reaction coordinate diagrams from of the barrier from the intermediate to native state in the absence (left) and in the presence (right) of G4. *Δ*G^‡^ is shown in black, *Δ*S^‡^ in cyan, and *Δ*H^‡^ in red.

**Table 1.**
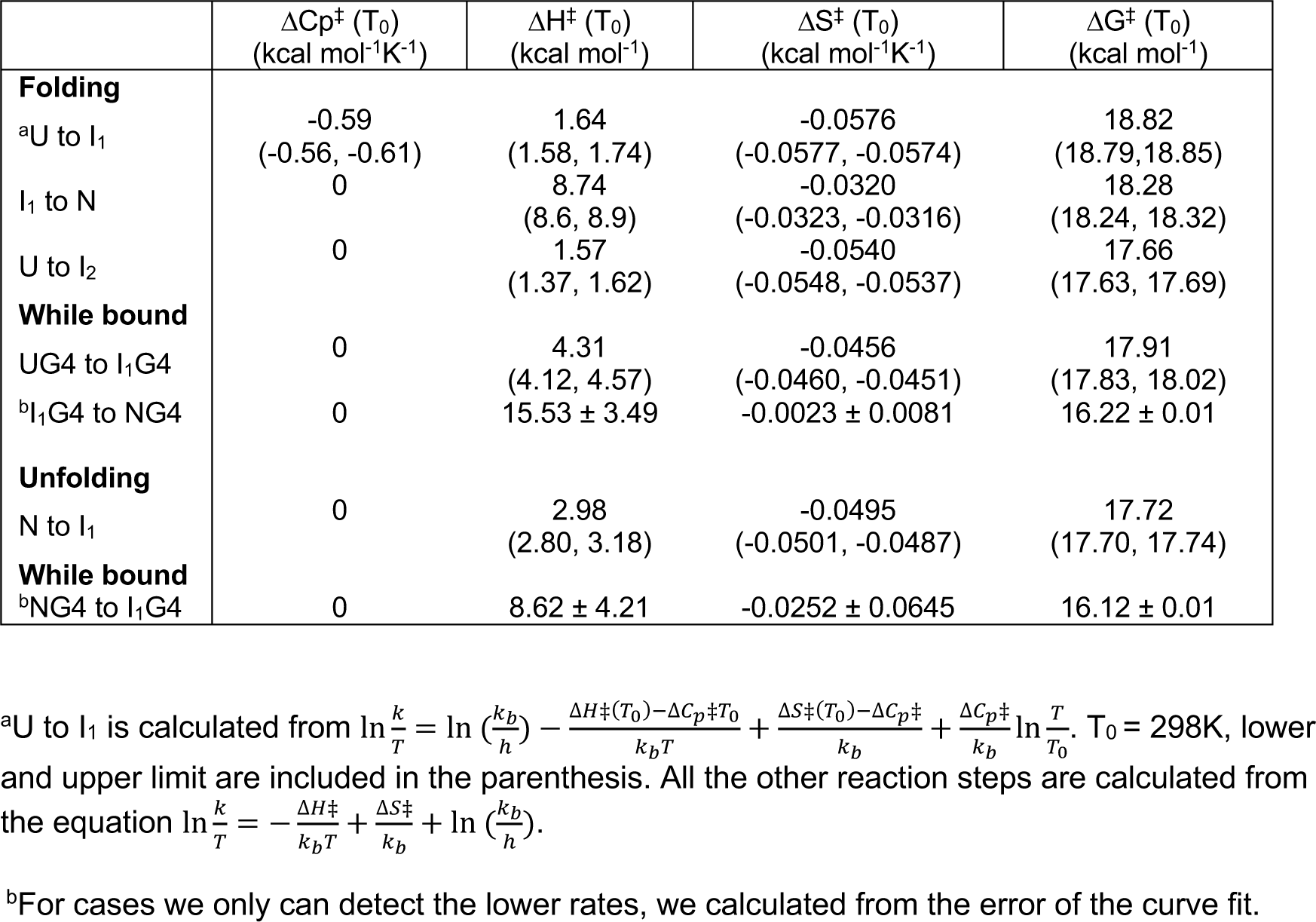
Activation parameters for the refolding and unfolding of TagRFP675 with and without G4. Calculated using mechanism from Fig. 1B, f and Fig. 2B. The errors of all parameters are reported using the lower and upper limits of the rates from the best fit.

## Discussion

In this study, we examined the effects of a G4 protein folding mechanism. Surprisingly, we found that the G4 catalyzed protein folding.

Acceleration of protein folding has primarily been posited as a unique function reserved for ATP-dependent chaperonin cages (2, 13, 14). Previous cases of folding while bound to ATP-independent chaperones have showed that while possible, folding was slower, not accelerated (3), with the rate of slowing depending on the strength of chaperone-protein interaction (15).

Although a previous case of folding acceleration by Arc repressor was observed with the addition of its DNA binding partner (16), this was attributed to increased dimerization speed due to reducing search space using 1D diffusion as opposed to folding catalysis (17).

The temperature-dependent experiments provide substantial clues as to the mechanism by which G4s catalyze protein folding. The increase in the enthalpy barriers with the addition of the G4 presumably arises from the formation of favorable contacts between the protein and the G4 that need to be broken for folding to continue. This observation is similar to the previous observation of proteins folding slower while bound to chaperone proteins (15).

The difference from previous observations with chaperone proteins in is that with the G4, offsetting thermodynamic forces ultimately cause an increase in rate constants. In the case of U to I_1_, there is a large change in heat capacity of the transition state when a G4 is present. Heat capacity has been previously shown to be related to changes in structure or in solvent-accessible surface area (18). Therefore, the effect of the G4 from the unfolded state contains two parts: 1) The catalysis of the U to I_1_ transition causes this step to outcompete the off-pathway U to I_2_ transition, leading to productive folding, and 2) this catalysis is due to changes in the structure or solvation of the U to I_1_ transition by the G4 (Fig 3B).

For I_1_ to N, the energetic driving forces are different. There is still an increase to the enthalpic barrier as for U to I_1_ presumably due to the need to break favorable interactions for folding to complete. However, for I_1_ to N, the G4 causes no discernable change in heat capacity but instead a large decrease in the entropy barrier (Fig 3C). Changes in entropy could be caused either by changes in solvation entropy or configurational entropy. However, the lack of change in heat capacity would suggest that the catalysis is more likely to be caused by changes in configurational entropy of the transition state. This observation is consistent with our previous observations of the importance of dynamics and oligomerization to the anti-aggregation activity of G4s (6, 8). A decrease in the configurational entropy barrier would suggest that the G4 increases number of possible pathways between the I_1_ and N states. To our knowledge, this is the first experimentally-derived analysis of the thermodynamic driving forces of protein folding catalysis by any chaperone.

Hsp60 chaperonins have been suggested to be required for folding by only 10% of the protein folding in *E. coli* and eukaryotic cells (19, 20), despite many proteins *in vitro* having slow folding rates. Proteins with slow folding speed like TagRFP (estimated folding time of tens of seconds or higher) are near the slow end of the folding time distribution of the proteome. These proteins may need additional help to speed up folding sufficiently to outpace degradation (21). The results here provide a potential explanation for why Hsp60 dependence of the proteome is relatively low, as folding acceleration of such slow folding proteins does not solely rely on chaperonins as previously thought. At the very least, folded G4s could also play this role, and it is possible that other macromolecules could as well. While the concentration of G4s in cells is not quantitatively known, they are known to increase as a function of stress (4). Moreover, the largest number of G4s in eukaryotes are likely found in ribosomal RNA extensions, creating a high local concentration of G4s at the site of original protein folding (22). As a result, the folding capacity of the cell could be much greater than previously characterized.

## Materials and Methods

### Expression and purification of TagRFP675

Wild type pBAD/HisD-TagRFP675 was transformed by heat shock into *E.coli* BL21(DE3) strain. TagRFP675 was inoculated at 37 °C for 3 hours at 700 rpm then was induced on Luria-Bertani (LB) agar ampicillin plates with 0.2% L-arabinose at 42 °C overnight. TagRFP675 cell pellet was collected with 170 mM NaCl. Purification of TagRFP675 is previously described (5). Final concentration and the purity of the protein were tested using both ThermoFisher NanoDrop UV-vis spectrophotometer and SDS-PAGE, respectively.

### Preparation of DNA G4 Seq 576

G4 DNA oligonucleotide Seq 576 5’-TGTCGGGCGGGGAGGGGGGG-3’ was purchased from Integrated DNA Technologies. Seq 576 was resuspended in filtered 10 mM potassium phosphate (pH 7.5) at room temperature prior to annealing. Annealing was performed by heating the sequence at 95 °C for 2 mins, then cooling from 95 °C to 25 °C at 1 °C per minute while shaking on Eppendorf ThermoMixer C. Seq 576 was then spun down by using VMR tabletop centrifuge before fluorescence kinetic experiments, with a stock concentration of Seq 576 of 100 μM.

### In vitro fluorescence kinetics experiments

Fluorescence intensities were monitored for a total 3-minute run at TagRFP675 native wavelength excitation/emission 598/675 nm, and near-native (fluorescent intermediate) wavelength excitation/emission 388/510 nm. All samples were introduced at 1 minute under constant stirring. Prior to sample injection, baseline fluorescence signal is 1.2 ± 0.3 at 598/675 nm, and 5.1 ± 0.7 at 388/510 nm. All experiments were performed at least duplicates for reproducibility. All kinetic experiments were performed on Agilent Cary Eclipse fluorescence spectrophotometer G9800A.

*Unfolding assay:* Freshly made 8M guanidine hydrochloride (GuHCl) was prepared in 10 mM potassium phosphate (pH 7.5) at room temperature. At 20 °C, GuHCl was diluted to 6M for all unfolding experiments in quartz fluorometer cuvettes. The final concentration of TagRFP675 were 0.5, 1 and 2 μM. And in the presence of G4, the final concentration of pre-incubated TagRFP675: Seq 576 mixture were 0.5:1, 1:2, and 2:4 μM.

*Refolding assay:* TagRFP675 was chemically denatured in 6M GuHCl overnight. At 20 °C, the final concentration of TagRFP675 were 0.25, 0.5 and 1 μM. In the presence of G4, Seq 576 was initially diluted in 10 mM phosphate buffer in the cuvette, then TagRFP675 was introduced at 1 minute while stirring. Final concentration of TagRFP675: Seq 576 mixture were 0.25:0.5, 0.5:1, and 1:2 μM. The final GuHCl concentrations in all the refolding assay were less than 50 mM. Of note, we have found that the amount of native state achieved in the experiment is dependent on the residual GuHCl level. Above assays were repeated at five other temperatures, 4, 25, 30, 37 and 40 °C. 10 mM potassium phosphate (pH 7.5) was pre-chilled on ice and pre-warmed in water bath for experiments at 4 °C and higher temperatures, respectively. For the topology dependence experiments only, methods were as previously described with higher residual GuHCl concentrations (5).

### Global fitting

All data were initially assessed to burst, single or double exponential using analytical functions by nonlinear regression in KinTek Explorer (KinTek Corporation) (23, 24). Goodness of fit was determined by chi^2^ (*X*^2^) divided by the degree of freedom (dof). Each thermodynamic cycle in the mechanism was constrained to a net free energy of 1. Mechanisms were built upon three-state model of protein folding which contains one on-path intermediate prior to the native state. For the folding of TagRFP675, output expressions at native and intermediate wavelength were defined by the native state and the intermediate states of TagRFP675. In the presence of quadruplex, TagRFP675-G4 complexes were included as contribution to the fluorescence signal at both wavelengths. All fitting were performed using the average from trials. There were no significant rate changes while comparing individual to the average trials. For data normalization, scaling factors were applied to each trace due to potential lamp fluctuation. The standard deviations of fluorescence background at both wavelengths were low compared to the *X*^2^ values used for model selection, ensuring models were not overfitted. Inner filter effect of fluorescence was auto corrected in Kintek Explorer. For well constrained rates in all experiment shown, the 95% confidence contour analysis was based on a calculated *X*^2^ threshold at boundary of 0.998 to allow estimating the upper and lower limits for each reaction rate. Rates above 0.998 were considered a good fit. Errors are reported using 0.99 as *X*^2^ threshold at boundary. By fitting only the refolding and not the unfolding kinetics using the mechanisms in Fig. 1B, *X*^2^/dof are a) 3.67, b) 3.06, c) 5.47, d) 4.18, e) 3.59, f) 1.45, suggesting model selection is not solely determined by unfolding kinetics. In fact, the best model would be the same even if only refolding data was considered. However, the likelihood of the best model is even higher compared to the other models (based on ratios of *X*^2^/dof values) when both folding and refolding data is considered.

Data were fit using Eyring equations (25) with and without heat capacity using SciPy.

### Predicting folding time from theory of protein kinetics

Folding speed (k_f_) of a protein can be predicted from chain length (N) as ln k_f_ = 16.15 – 1.28N^0.5^ using an earlier theory by Thirumalai (11) fitted to a data set of 80 proteins with experimentally measured folding speed (26). This formulation estimates TagRFP folding speed to be 22 seconds. An alternate approach by Plaxco, Baker and Simons predicts folding speed using contact order calculated from the structure (12). Using this formalism trained against the folding speed data (27) predicts folding time of TagRFP to be 18 seconds. A slight variation of the contact order theory taking care of higher order contacts predicts TagRFP folding speed to be 38 seconds.

### Estimating folding time from data

An effective folding time of TagRFP from the experimental data in the absence of G4 can be estimated to compare against the theoretical predictions. Since theoretical predictions do not account for off pathway intermediates, we consider only the on-pathway intermediate (I_1_) and ignore I_2_ for this comparison. Ignoring I_2_, folding proceeds in two consecutive steps: first transitioning to the intermediate (I_1_) with an average folding time of 12.5 seconds (using a rate of 0.08 s^-1^ at 20 °C reported in Table S1) followed by a second transition from I_1_ to the native state with an average time of 4.7 seconds (using a rate of 0.21 s^-1^ at 20 °C reported in Table S1). Thus, the total average folding time is approximately 17 seconds. If the full network with off-pathway intermediate is considered, the folding time would be even slower.

## Acknowledgments

The authors would like to thank S. Bromberg for making the folding energy landscape. This work was funded by NIH R35GM142442 (SH), R01GM138901 (KG), NSF 2236541 (FS).

---

**Fig. S1.**
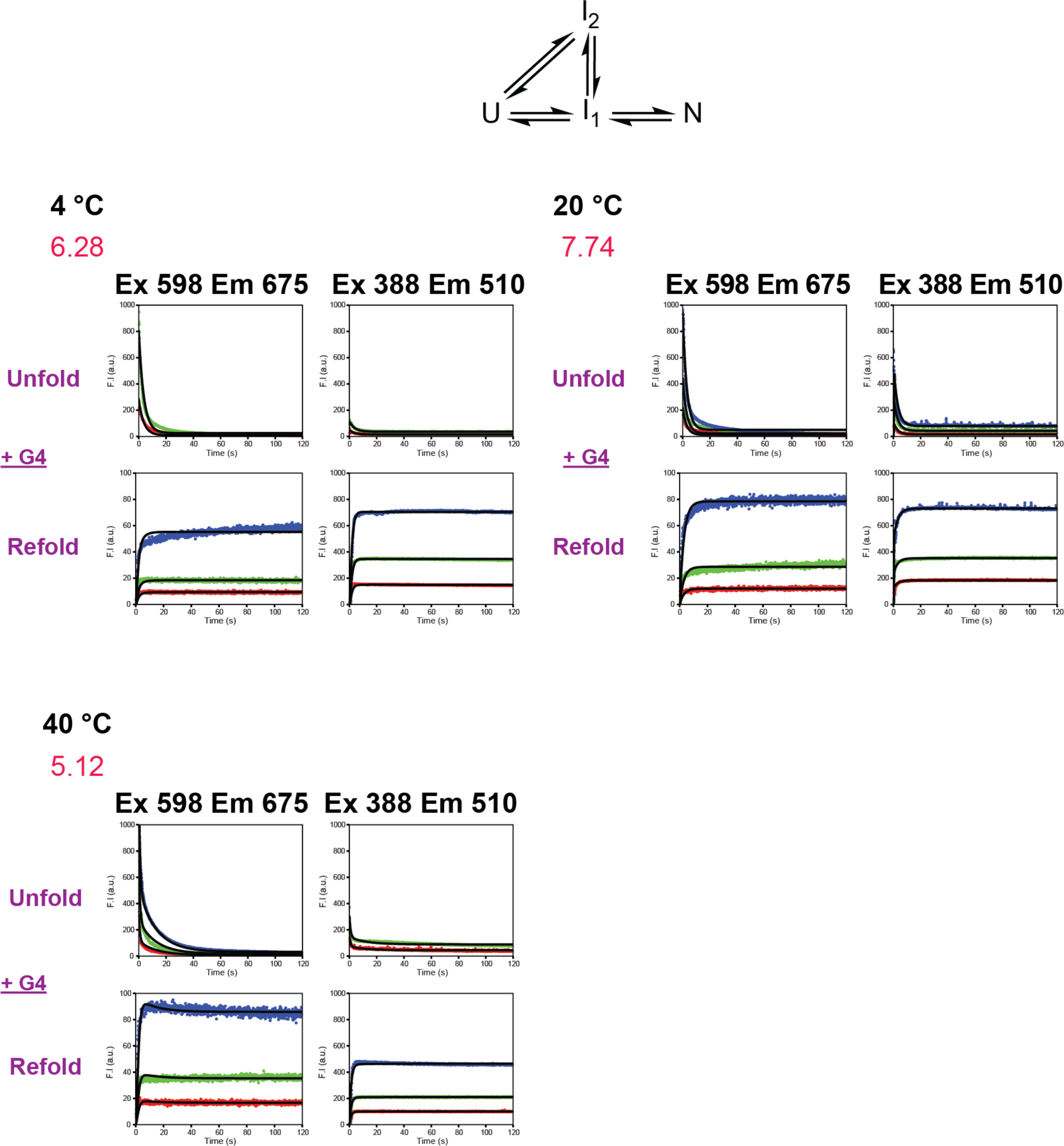
Control fits at 4, 20 and 40 °C. Global fitting of TagRFP675 with G4 using best fit mechanism from Fig. 2B, f) (top). Poor *X*^2^/dof values (labeled in red) at all temperatures indicate that G4 must be accounted for in the folding mechanism when G4 is present in the experiment.

**Fig. S2.**
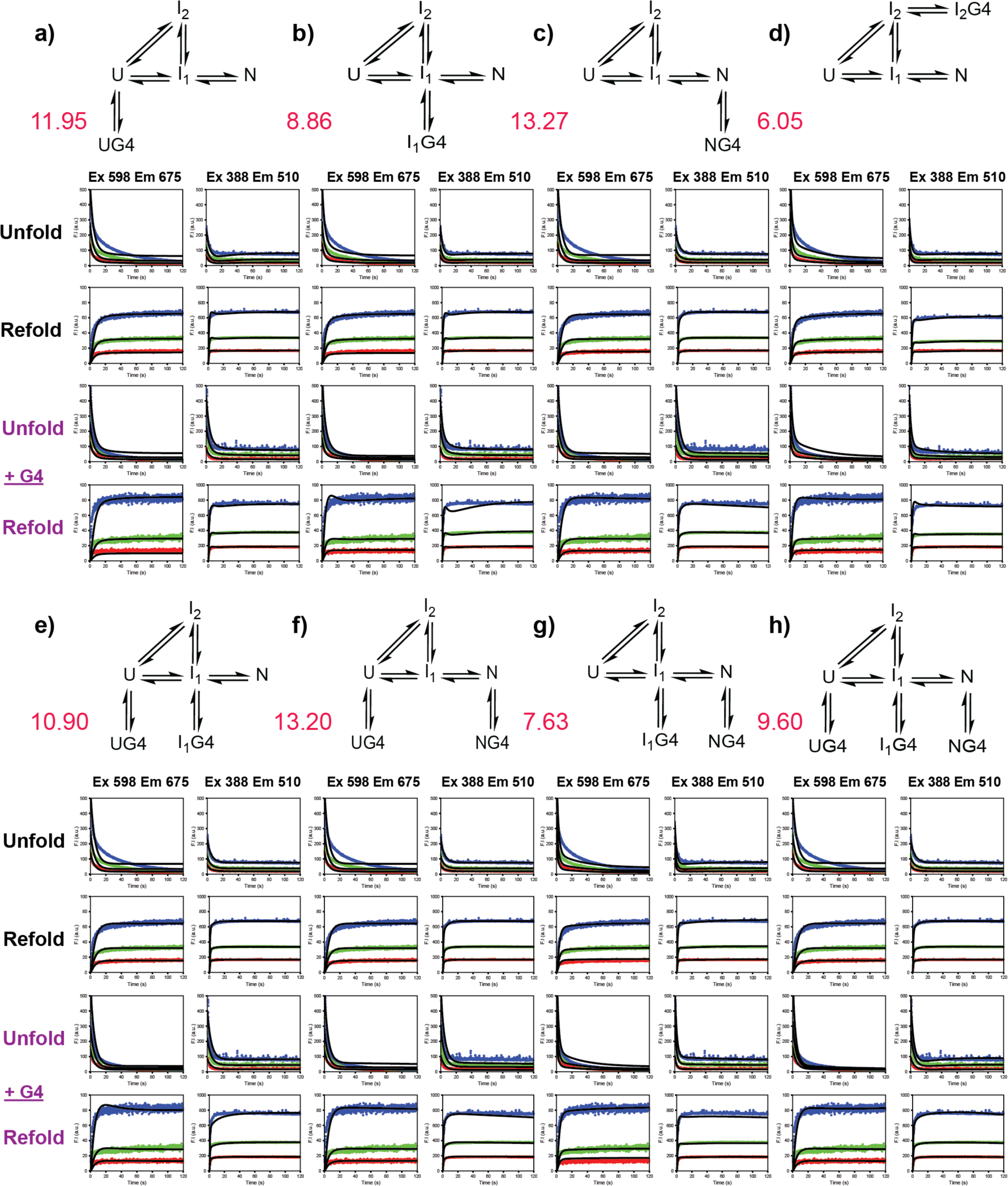
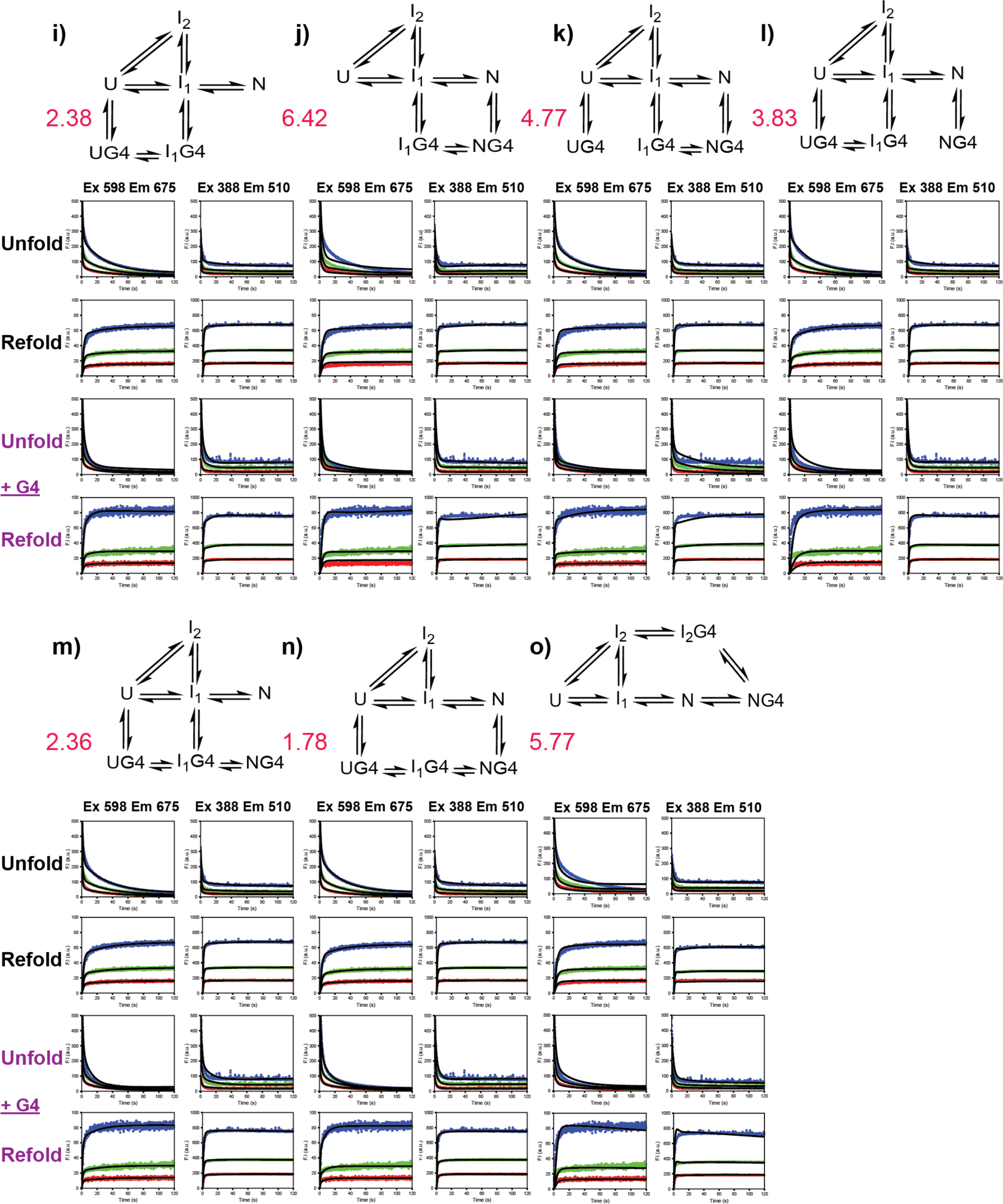
Systematic test of mechanism for folding of TagRFP675 bound in the presence of G4 at 20 °C. a-h). mechanisms in which no folding while bound occurs, i-o). folding while bound mechanisms. Smaller *X*^2^/dof values (labeled in red) indicate that the protein folds while bound to the G4.

**Fig. S3.**
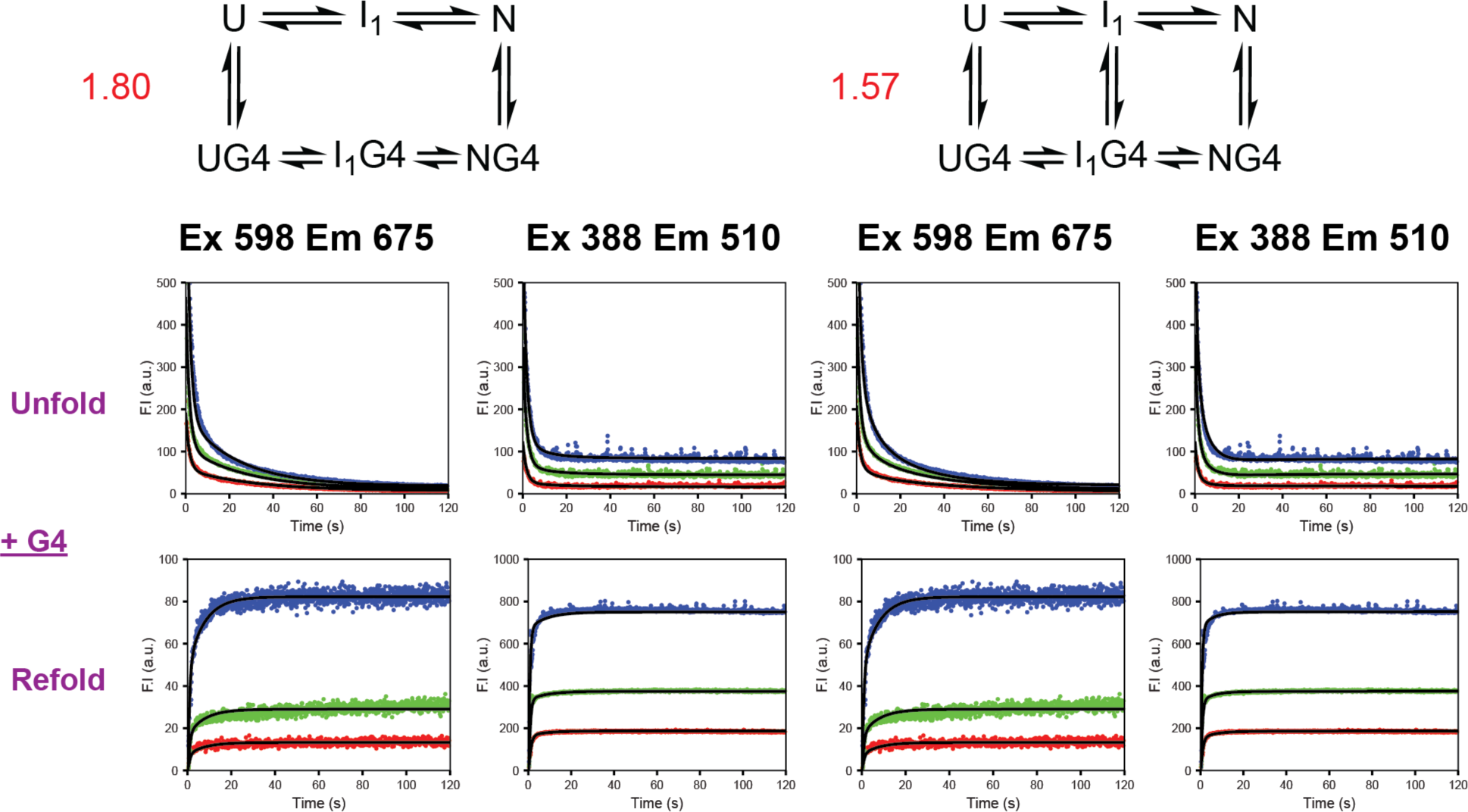
G4-assisting mechanism at 20 °C. Off-pathway intermediate I_2_ is no longer required with G4 presence. The mechanism on the right contains an intermediate binding step, which does not drastically improve the fit. Mechanism on the left is the minimum model.

**Fig. S4.**
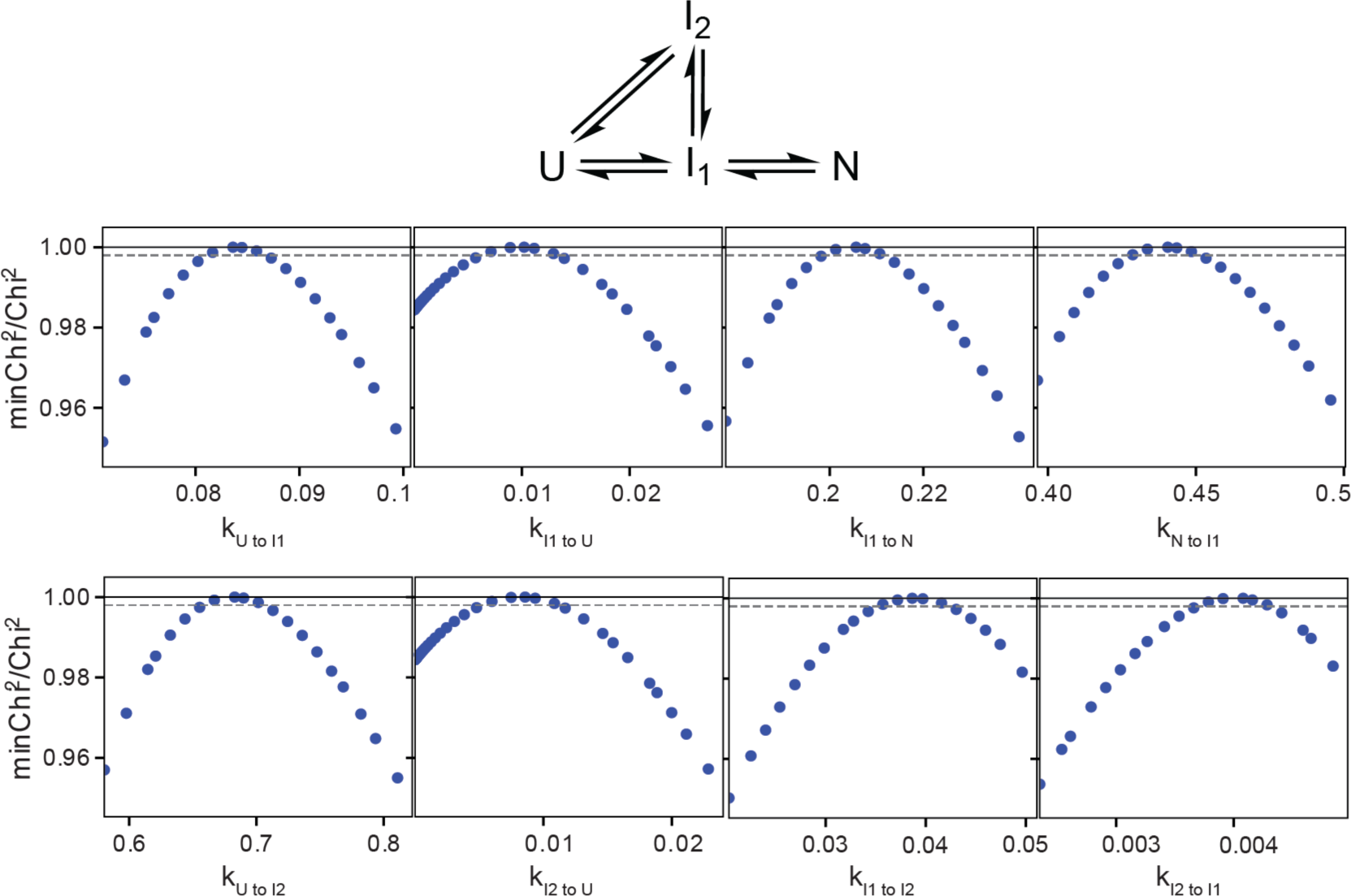
95% confidence contour analysis at 20 °C. 1D confidence contour analysis calculated from mechanism in Fig. 1B, f, with the *X*^2^ threshold limit set at 0.95. Y-axis (minChi^2^/Chi^2^) represents Z factor. *X*^2^ threshold at boundary sets at 0.998 (dashed grey line) from global fitting providing lower and upper limits for each reaction.

**Fig. S5.**
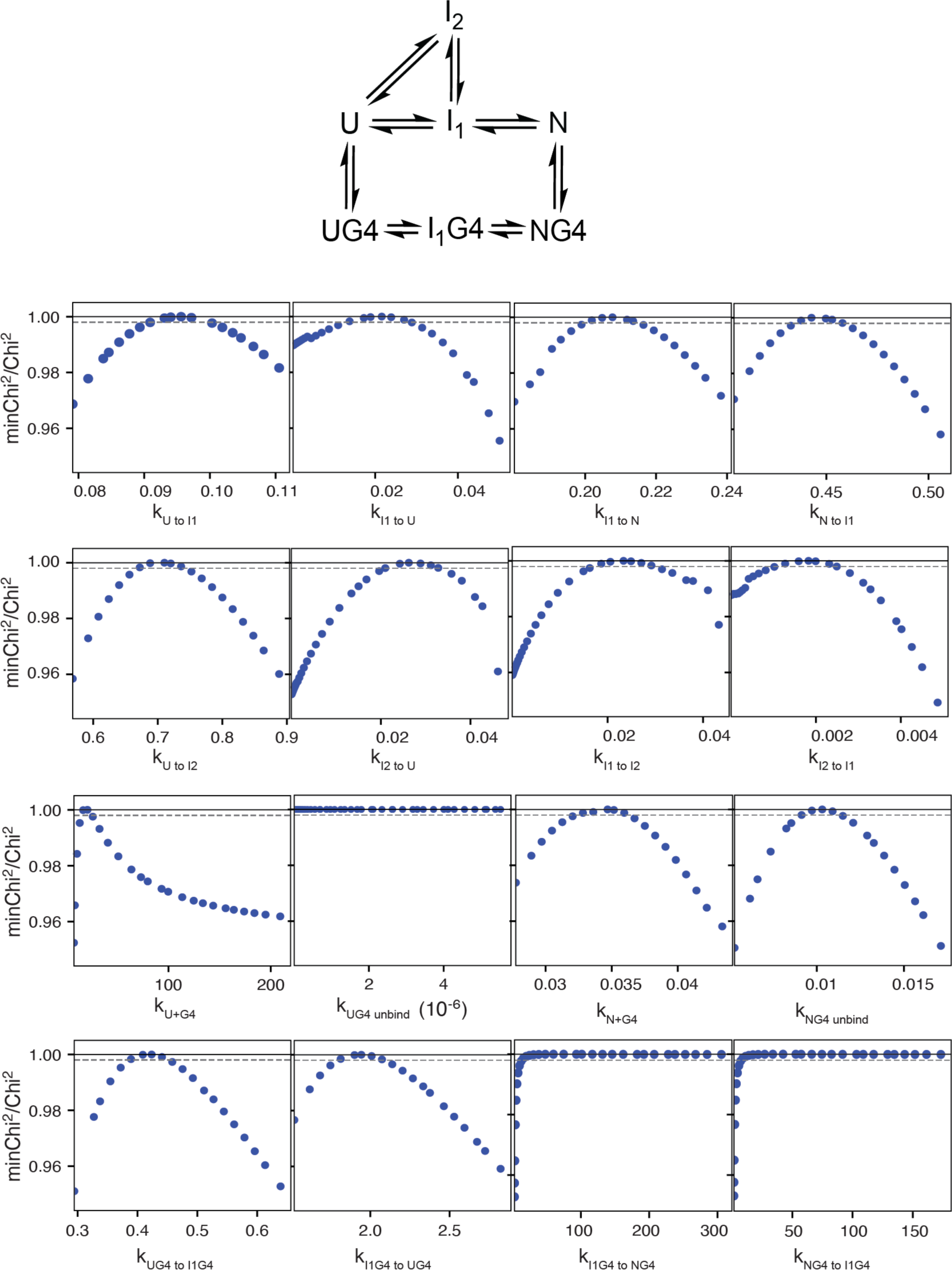
95% confidence contour analysis with G4 at 20 °C. 1D confidence contour analysis of mechanism in Fig. 2B, with the *X*^2^ threshold limit sets at 0.95. In several cases, only lower limits can be defined. The lower bound for k_I1G4 to NG4_ is 6.41 s^-1^ and k_NG4 to I1G4_ is 3.63 s^-1^. Above the lower limit, the data defines only the ratio of forward and reverse of I_1_G4 and NG4.

**Fig. S6.**
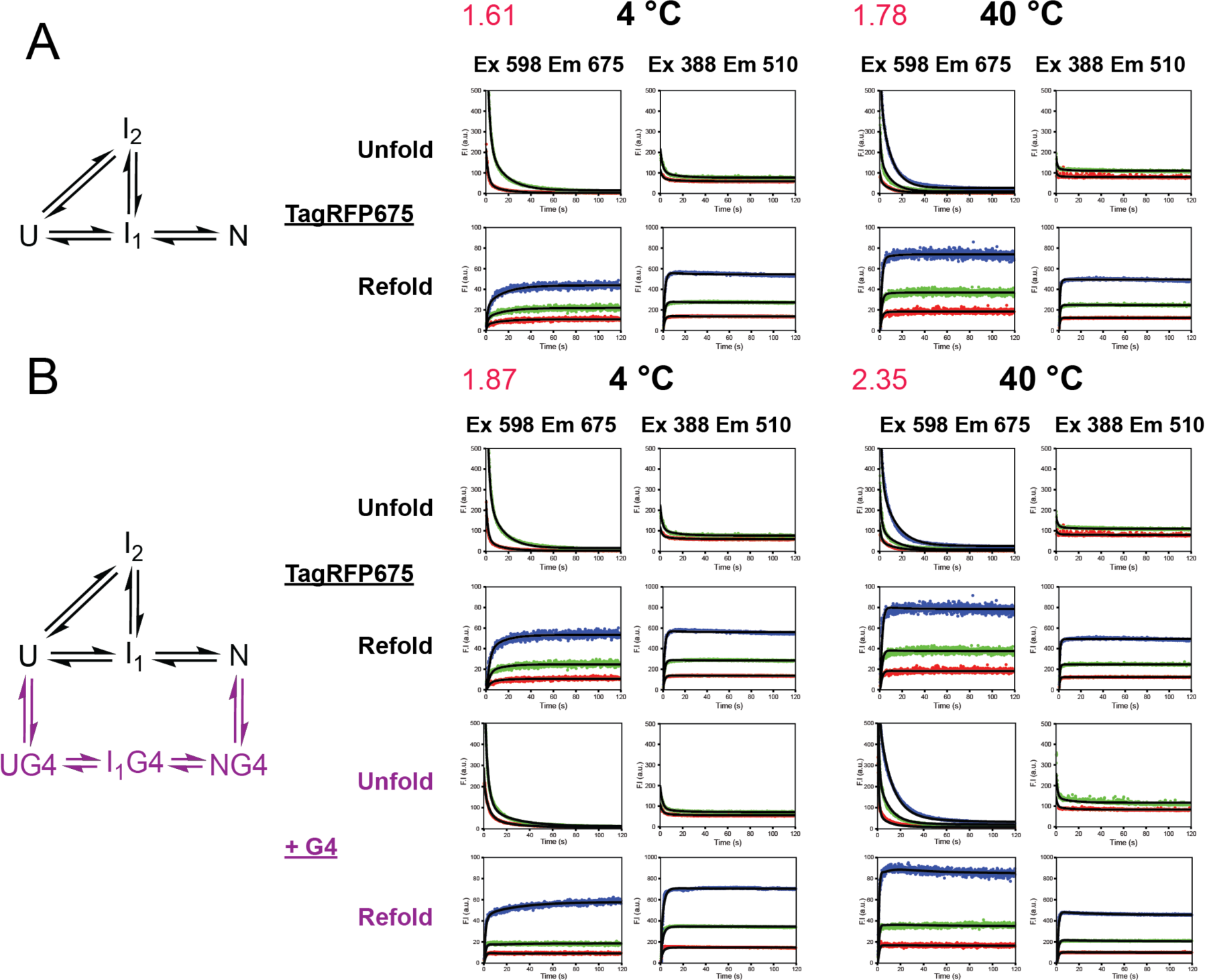
Examples of mechanisms. A. TagRFP675 folding by itself at 4 and 40 °C. B. TagRFP675 in the presence of G4 at 4 and 40 °C. Data was best fit using the same mechanism in Fig. 1B(f) and Fig. 2B respectively. *X*^2^/dof are labeled in red. All are 2:1 ratio of G4:TagRFP675. For unfolding experiments, TagRFP675 concentrations are 0.25 (red) and 0.5 (green) μM at 4°C. 0.5 (red), 1 (green), and 2 (blue) μM at native wavelength, 0.5 (red) and 1 (green) μM at intermediate wavelength at 40 °C. For refolding experiments, TagRFP675 concentrations are 0.25 (red), 0.5 (green), and 1 (blue) μM at both temperatures.

**Figure S7.**
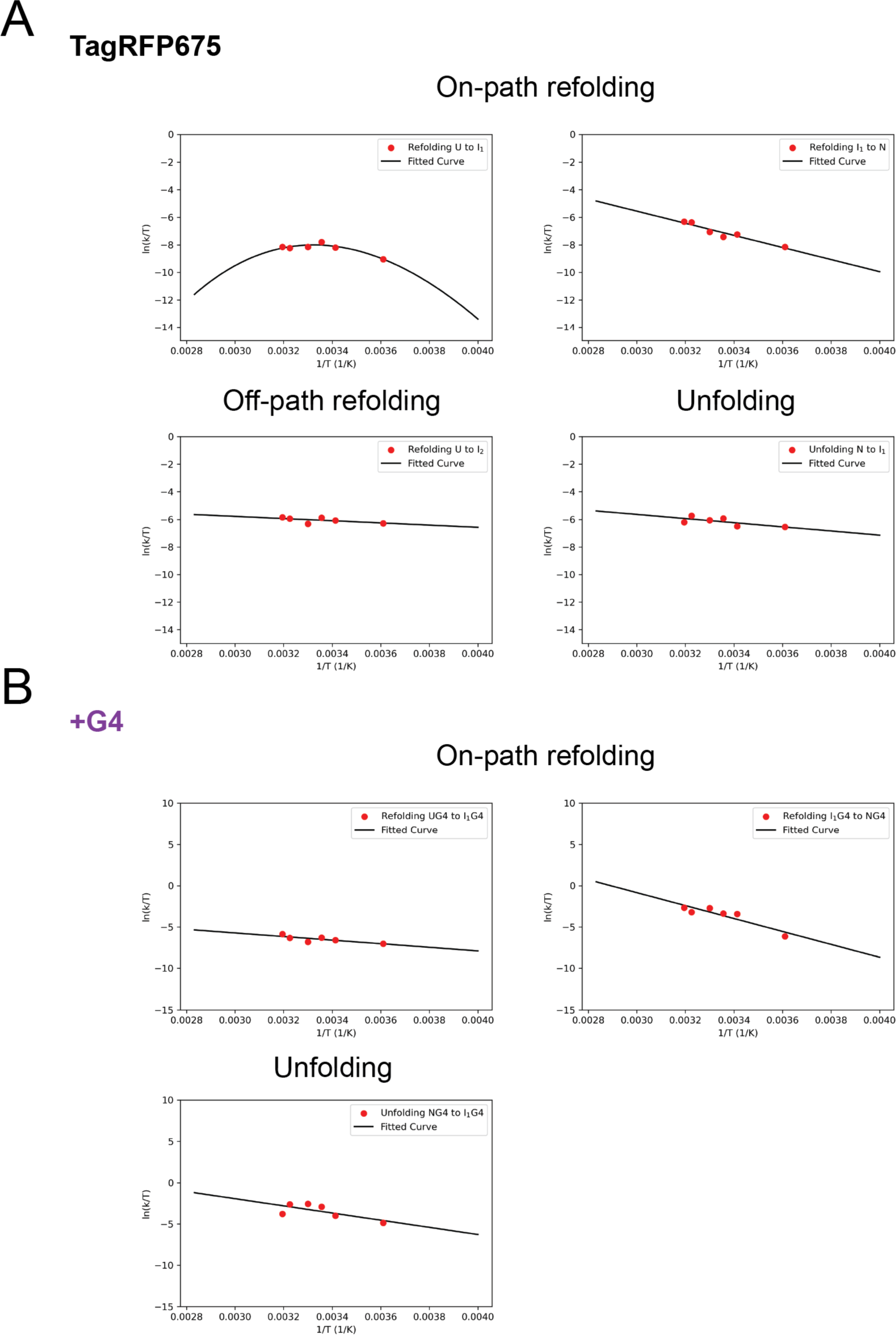
Eyring plots. For A. TagRFP675 folding and unfolding alone and B. for TagRFP675 folding and unfolding with G4. Curvature was only observed for step U to I_1_.

**Fig. S8.**
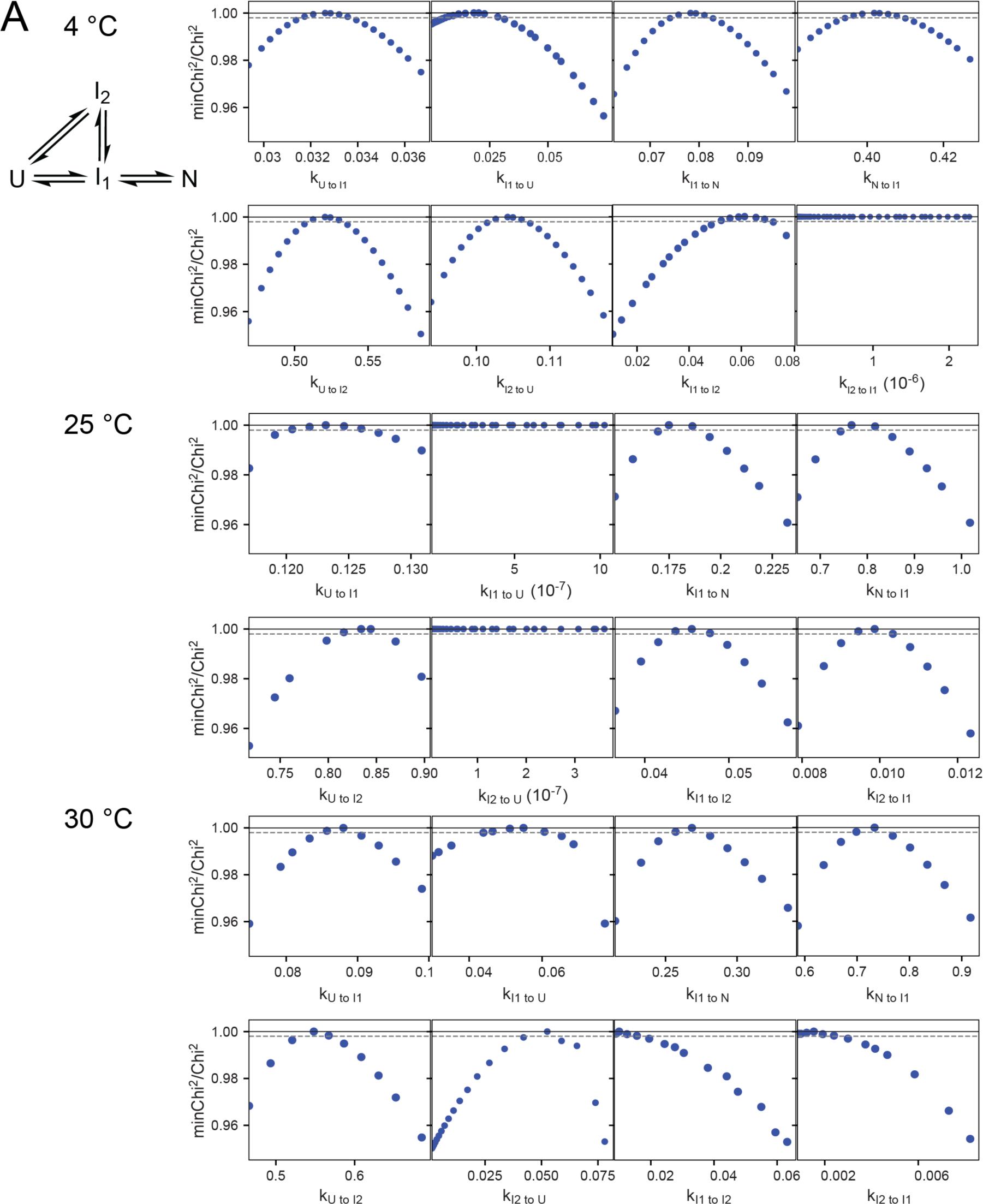
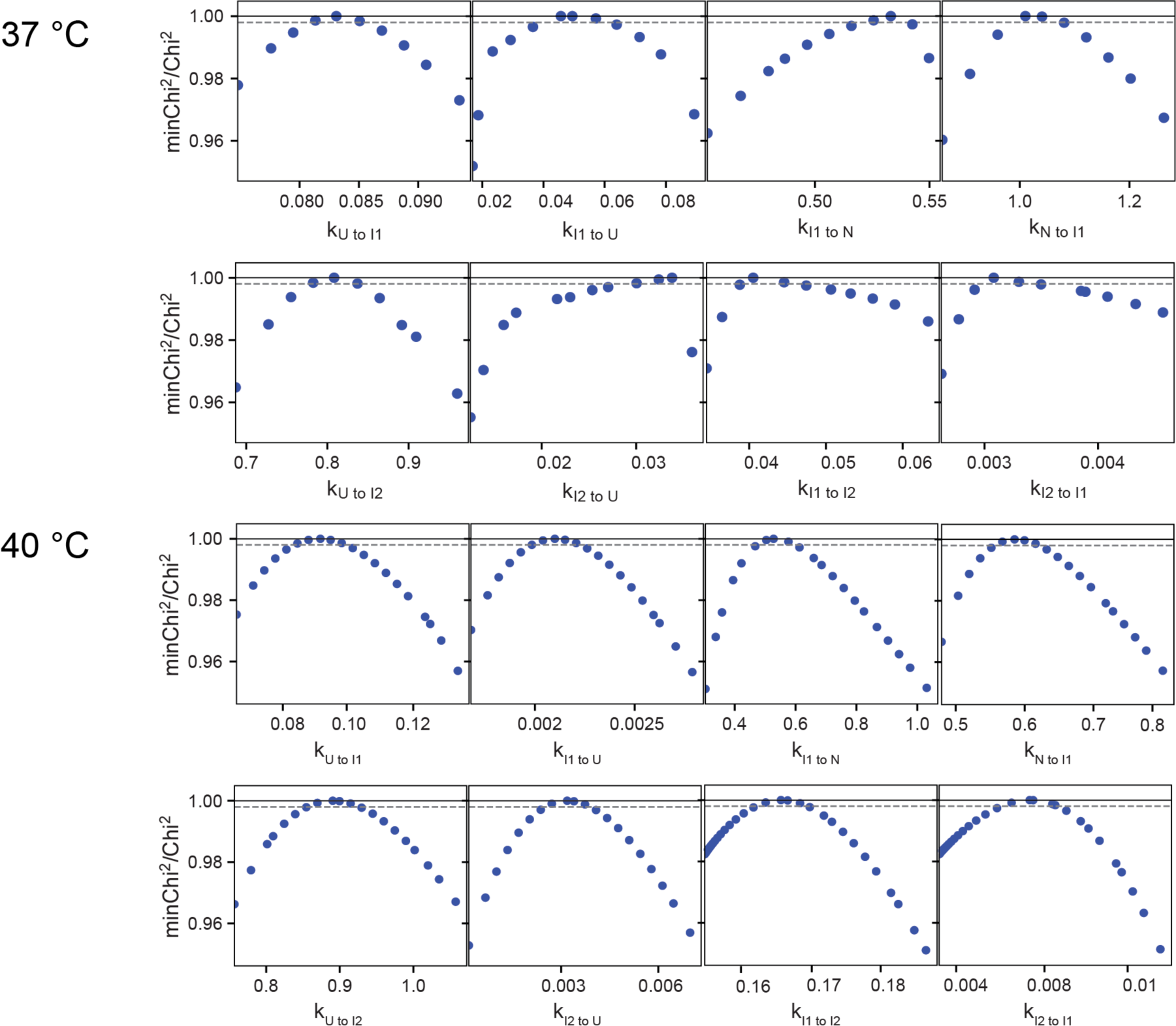
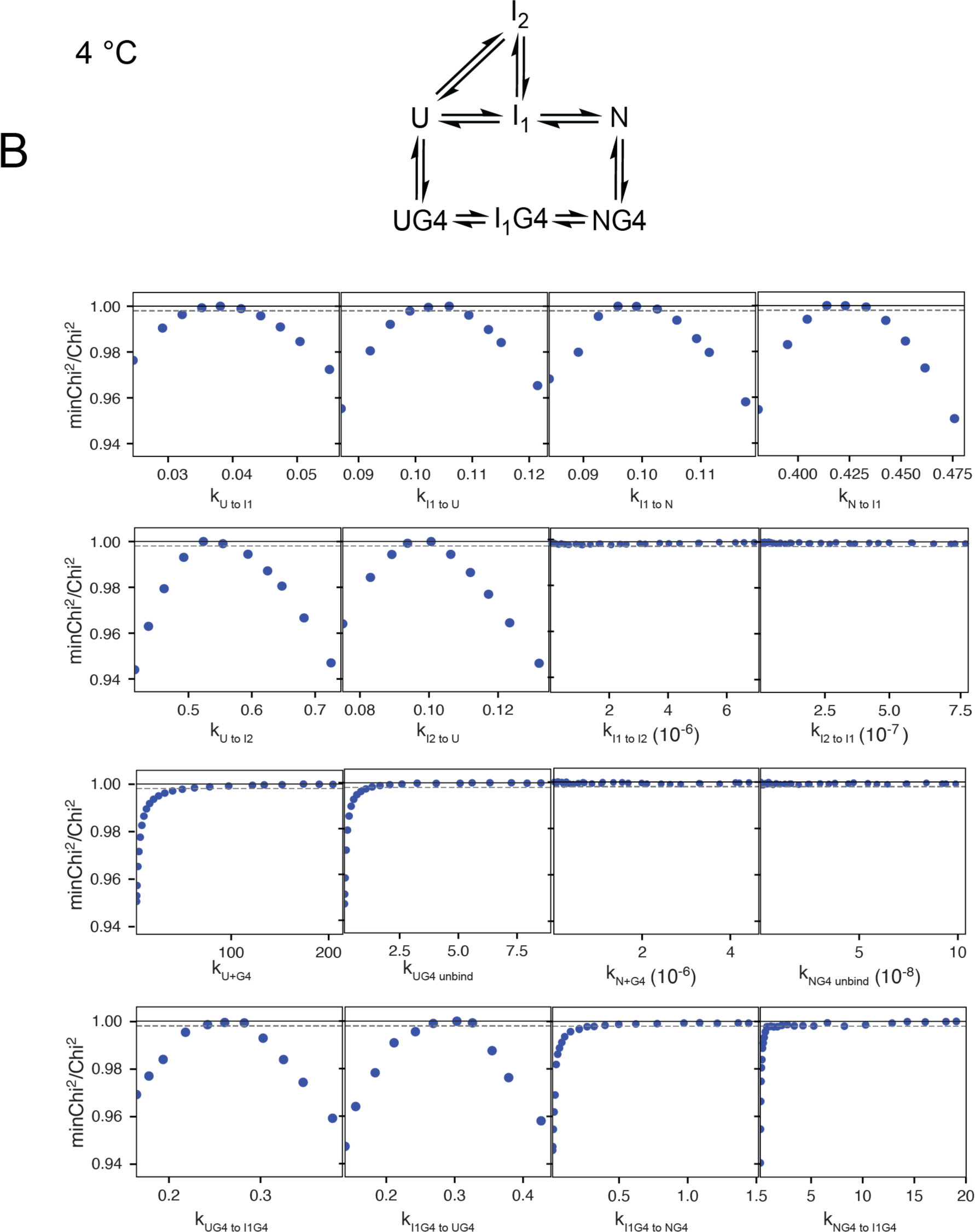
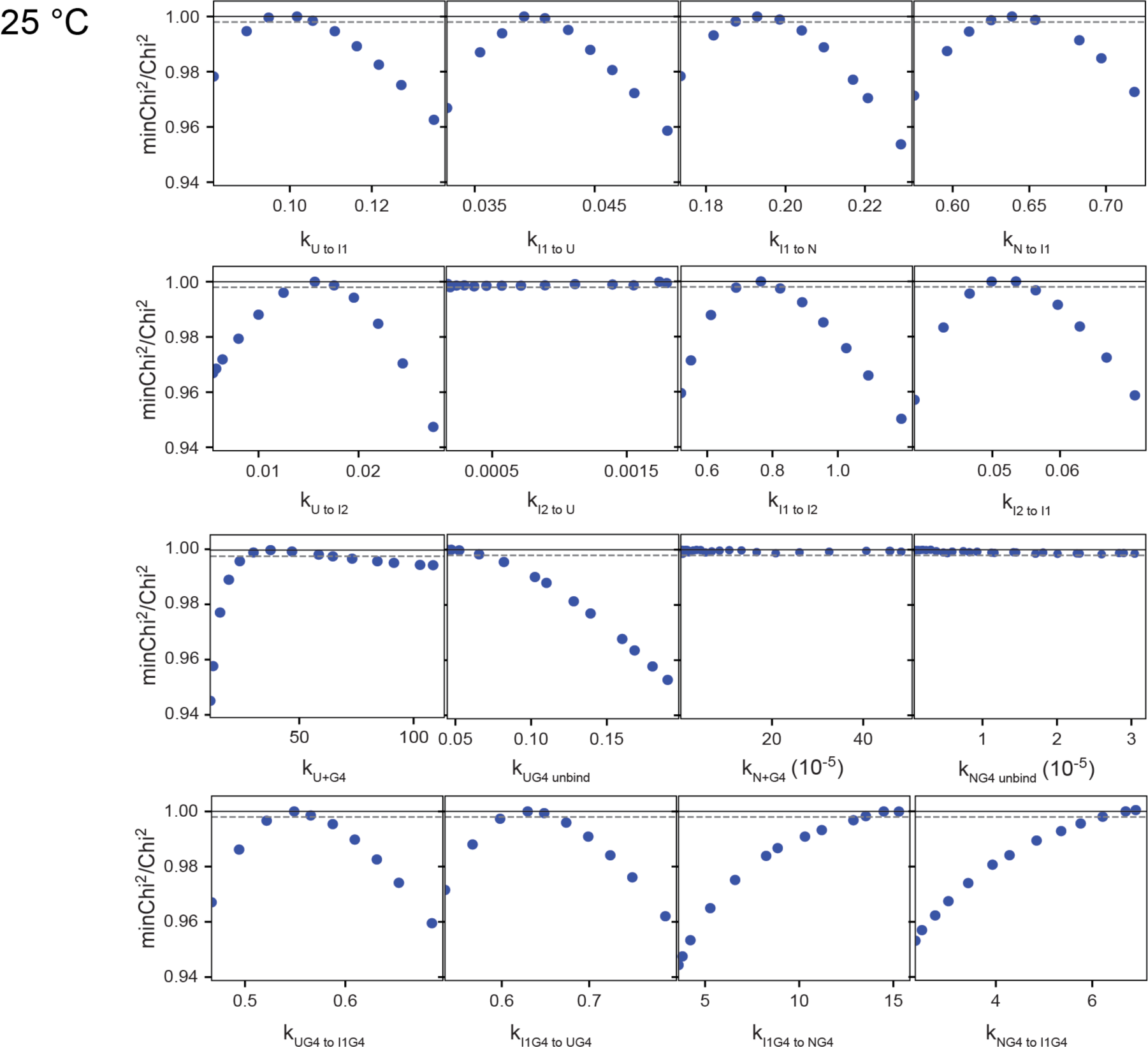
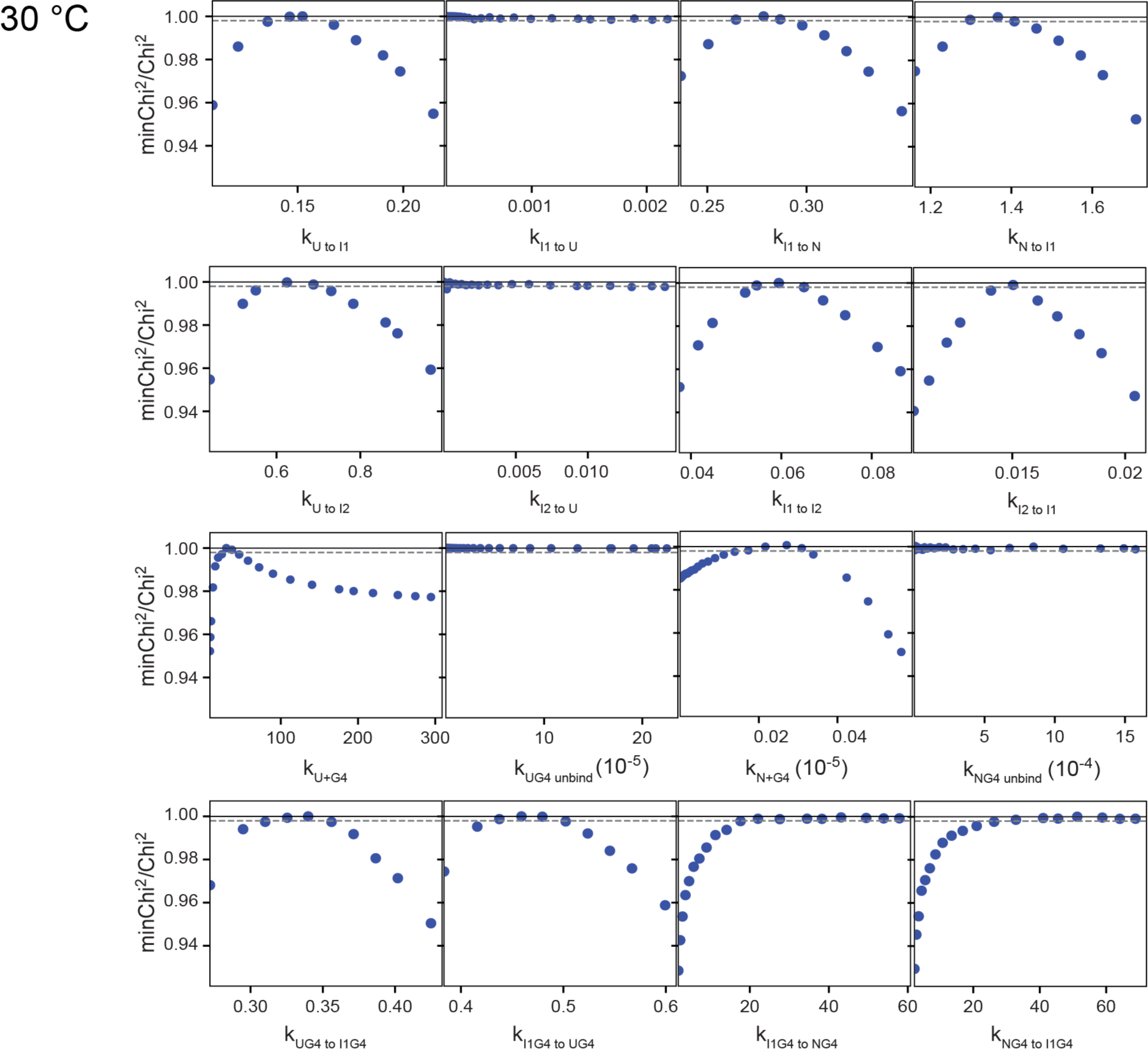
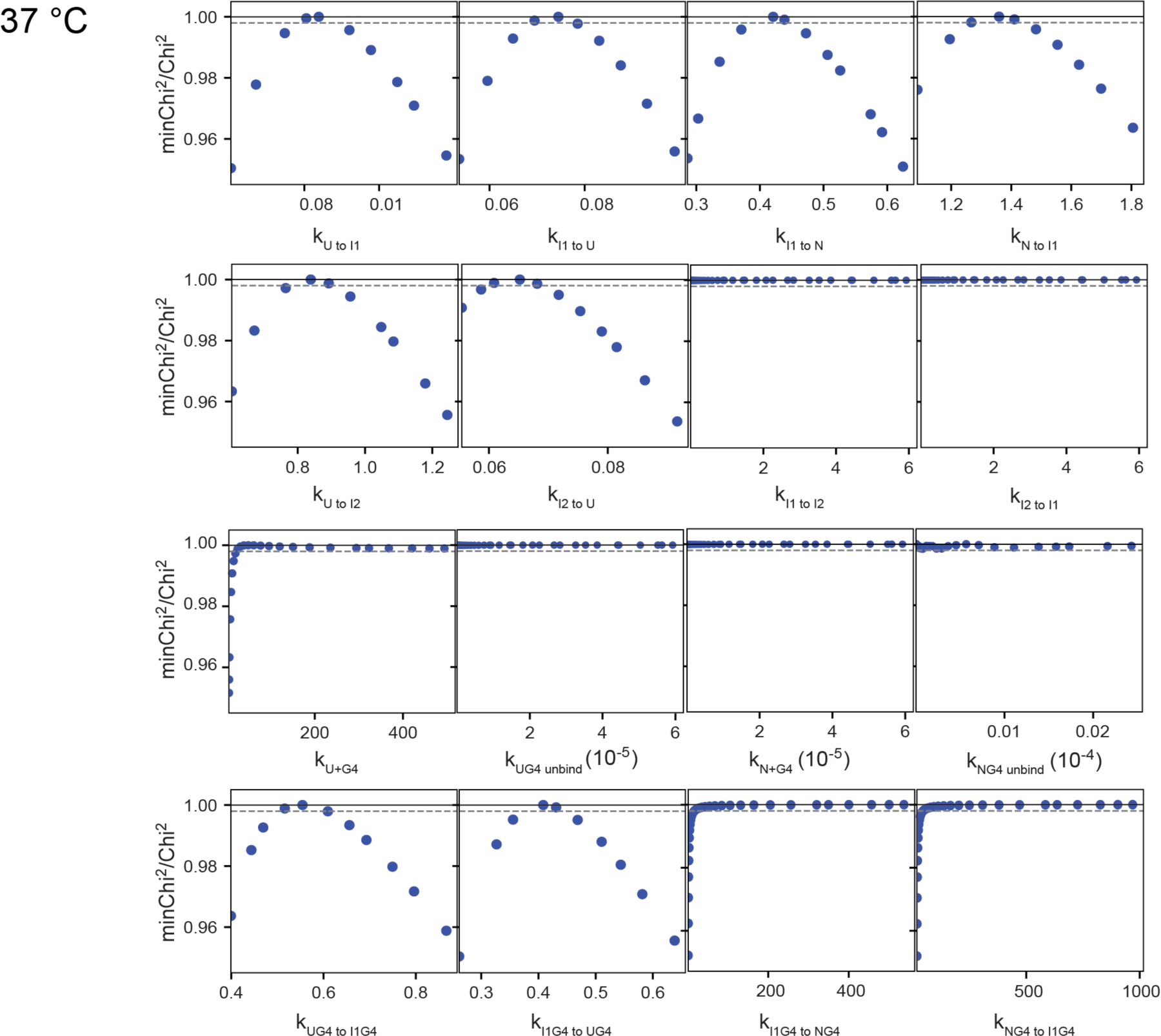
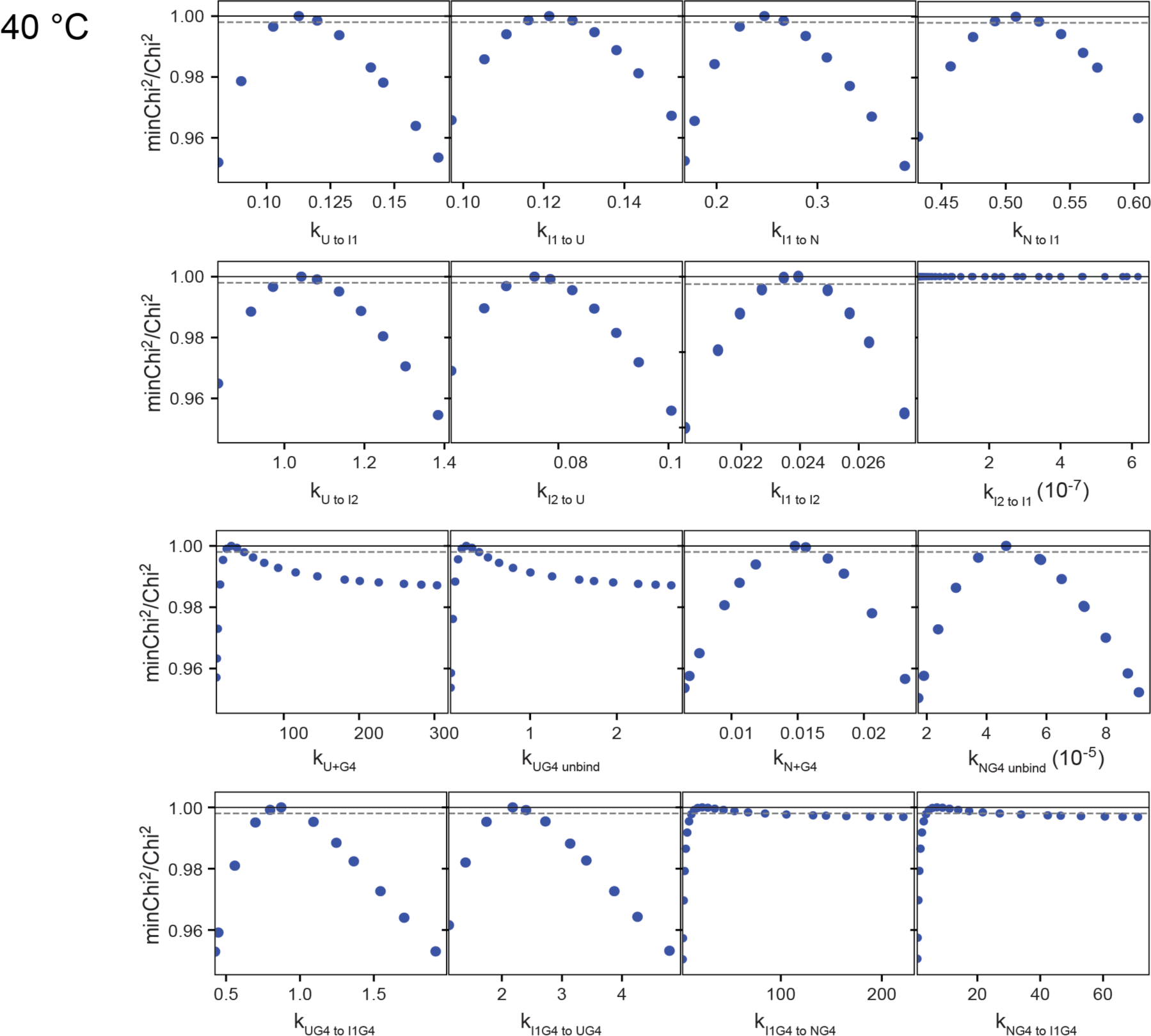
95% confidence contour analysis. A. TagRFP675 from 4 to 40 °C. *X*^2^ at 4, 25, 30, 37, 40 °C are 1.61, 1.29, 1.32, 1.41, and 1.78, respectively. B. with G4 from 4 to 40 °C. *X*^2^ at 4, 25, 30, 37, 40 °C are 1.87, 1.56, 1.73, 3.64, and 2.35 respectively. The major contributing error of the fit at 37 °C is from the unfolding step I_1_G4 to UG4, which we therefore did not quantify as a function of temperature. The *X*^2^ threshold limit was set at 0.95. Only lower limits can be defined for several steps.

**Table S1.** Rate constants for TagRFP675 at 4, 20, 25, 30, 37 and 40 °C without G4.

| **k (s<sup>-1</sup>)** | **4 °C** | **lower,**<br>**upper limit** | **20 °C** | **lower,**<br>**upper limit** | **25 °C** | **lower,**<br>**upper limit** |
| --- | --- | --- | --- | --- | --- | --- |
| **Folding** |  |  |  |  |  |  |
| U to I<sub>1</sub> | 0.033 ± 0.003 | (0.030, 0.036) | 0.08 ± 0.01 | (0.07, 0.09) | 0.124 ± 0.002 | (0.12, 0.126) |
| I<sub>1</sub> to N | 0.08 ± 0.01 | (0.07, 0.09) | 0.21 ± 0.01 | (0.2, 0.22) | 0.18 ± 0.006 | (0.174, 0.186) |
| U to I<sub>2</sub> | 0.52 ± 0.02 | (0.50, 0.54) | 0.68 ± 0.03 | (0.65, 0.72) | 0.84 ± 0.02 | (0.82, 0.86) |
| I<sub>1</sub> to I<sub>2</sub> | < 0.04 | (0, 0.04) | 0.04 ± 0.01 | (0.03, 0.05) | 0.046 ± 0.002 | (0.044, 0.048) |
| **Unfolding** |  |  |  |  |  |  |
| I<sub>1</sub> to U | <0.1 | ns | <0.1 | ns | <0.1 | ns |
| N to I<sub>1</sub> | 0.40 ± 0.01 | (0.39, 0.41) | 0.44 ± 0.01 | (0.43, 0.45) | 0.79 ± 0.03 | (0.76, 0.82) |
| I<sub>2</sub> to U | 0.105 ± 0.005 | (0.10, 0.11) | <0.1 | ns | <0.1 | ns |
| I<sub>2</sub> to I<sub>1</sub> | <0.1 | ns | <0.1 | ns | <0.1 | (0.009, 0.01) |

| **k (s<sup>-1</sup>)** | **30 °C** | **lower,**<br>**upper limit** | **37 °C** | **lower,**<br>**upper limit** | **40 °C** | **lower,**<br>**upper limit** |
| --- | --- | --- | --- | --- | --- | --- |
| **Folding** |  |  |  |  |  |  |
| U to I<sub>1</sub> | 0.088 ± 0.002 | (0.086, 0.09) | 0.083 ± 0.002 | (0.081, 0.085) | 0.09 ± 0.01 | (0.08, 0.1) |
| I<sub>1</sub> to N | 0.26 ± 0.01 | (0.25, 0.27) | 0.535 ± 0.015 | (0.52, 0.55) | 0.57 ± 0.08 | (0.49, 0.65) |
| U to I<sub>2</sub> | 0.55 ± 0.02 | (0.53, 0.57) | 0.81 ± 0.03 | (0.78, 0.84) | 0.9 ± 0.1 | (0.8, 1) |
| I<sub>1</sub> to I<sub>2</sub> | < 0.02 | (0, 0.02) | < 0.04 | (0.02, 0.04) | 0.17 ± 0.01 | (0.16, 0.18) |
| **Unfolding** |  |  |  |  |  |  |
| I<sub>1</sub> to U | 0.05 ± 0.01 | (0.04, 0.06) | 0.05 ± 0.01 | (0.04, 0.06) | <0.1 | ns |
| N to I<sub>1</sub> | 0.71 ± 0.01 | (0.7, 0.73) | 1.01 ± 0.02 | (0.99, 1.04) | 0.63 ± 0.06 | (0.57, 0.69) |
| I<sub>2</sub> to U | <0.1 | (0.04, 0.06) | <0.1 | (0.03, 0.04) | <0.1 | ns |
| I<sub>2</sub> to I<sub>1</sub> | <0.1 | (0, 0.003) | <0.1 | (0.002, 0.004) | <0.1 | ns |

**Table S2.** Rate constants for TagRFP675 with G4 at 4, 20, 25, 30, 37 and 40 °C.

| **k** | **4 °C** | **lower,**<br>**upper limit** | **20 °C** | **lower,**<br>**upper limit** | **25 °C** | **lower,**<br>**upper limit** |
| --- | --- | --- | --- | --- | --- | --- |
| **Folding** |  |  |  |  |  |  |
| U to I<sub>1</sub> | 0.04 ± 0.01 | (0.03, 0.05) | 0.09 ± 0.01 | (0.08, 0.1) | 0.101 ± 0.001 | (0.1, 0.102) |
| I<sub>1</sub> to N | 0.09 ± 0.01 | (0.08, 0.1) | 0.21 ± 0.01 | (0.2, 0.22) | 0.196 ± 0.003 | (0.193, 0.199) |
| U to I<sub>2</sub> | 0.56 ± 0.06 | (0.50, 0.62) | 0.73 ± 0.06 | (0.67, 0.79) | 0.77 ± 0.04 | (0.73, 0.81) |
| I<sub>1</sub> to I<sub>2</sub> | ns | ns | 0.02 ± 0.01 | (0.01, 0.03) | 0.01 ± 0.01 | (0, 0.02) |
| **While bound** |  |  |  |  |  |  |
| UG4 to I<sub>1</sub>G4 | 0.25 ± 0.05 | (0.2, 0.3) | 0.4 ± 0.1 | (0.3, 0.5) | 0.55 ± 0.05 | (0.5, 0.6) |
| I<sub>1</sub>G4 to NG4 | 0.6 | (0.5, 6) | 9.56 | (6.41, 306) | 10.3 | (10.3, 14.5) |
| **Unfolding** |  |  |  |  |  |  |
| I<sub>1</sub> to U | 0.1 | (0.1, 0.11) | 0.02 | (0, 0.04) | 0.035 ± 0.005 | (0.03, 0.04) |
| N to I<sub>1</sub> | 0.45 ± 0.05 | (0.4, 0.5) | 0.44 ± 0.02 | (0.42, 0.46) | 0.635 ± 0.005 | (0.63, 0.64) |
| I<sub>2</sub> to U | 0.1 | (0.09, 0.1) | 0.021 | (0, 0.04) | 0.05 ± 0.01 | (0.04, 0.06) |
| I<sub>2</sub> to I<sub>1</sub> | ns | ns | ns | ns | ns | ns |
| **While bound** |  |  |  |  |  |  |
| NG4 to I<sub>1</sub>G4 | 2.15 | (2, 19.1) | 5.44 | (3.63, 173) | 6.69 | (6.6, 9.4) |
| I<sub>1</sub>G4 to UG4 | 0.25 ± 0.05 | (0.2, 0.3) | 1.7 | (1.6, 2.2) | 0.63 | (0.62, 0.64) |
| **Binding** |  |  |  |  |  |  |
| U+G4 | 20.4 | (19, 519) | 23 | (13, 204) | 39.3 | (24, 109) |
| N+G4 | ns | ns | 0.035 | (0.03, 0.04) | ns | ns |
| **Dissociation** |  |  |  |  |  |  |
| UG4 | ns | ns | ns | ns | ns | ns |
| NG4 | ns | ns | ns | ns | ns | ns |

Table S2 continued
| k | 30 °C | lower,<br>upper limit | 37 °C | lower, upper<br>limit | 40 °C | lower,<br>upper limit |
| --- | --- | --- | --- | --- | --- | --- |
| **Folding** |  |  |  |  |  |  |
| U to I<sub>1</sub> | 0.147 ± 0.002 | (0.145, 0.149) | 0.08 ± 0.01 | (0.07, 0.09) | 0.11 ± 0.1 | (0.10, 0.12) |
| I<sub>1</sub> to N | 0.277 ± 0.001 | (0.276, 0.278) | 0.43 ± 0.06 | (0.37, 0.49) | 0.25 ± 0.03 | (0.22, 0.28) |
| U to I<sub>2</sub> | 0.65 ± 0.01 | (0.64, 0.66) | 0.85 ± 0.10 | (0.75, 0.95) | 1.06 ± 0.09 | (0.97, 1.15) |
| I<sub>1</sub> to I<sub>2</sub> | 0.065 ± 0.005 | (0.06, 0.07) | ns | ns | 0.024 | (0.022, 0.026) |
| **While bound** |  |  |  |  |  |  |
| UG4 to I<sub>1</sub>G4 | 0.34 ± 0.01 | (0.33, 0.35) | 0.56 ± 0.09 | (0.47, 0.65) | 0.9 ± 0.2 | (0.7, 1.1) |
| I<sub>1</sub>G4 to NG4 | 20 | (9.11, 57.9) | 12.7 | (4.61, 536) | 22.1 | (7.25, 221) |
| **Unfolding** |  |  |  |  |  |  |
| I<sub>1</sub> to U | ns | ns | 0.074 ± 0.09 | (0.065, 0.083) | 0.12 ± 0.01 | (0.11, 0.13) |
| N to I<sub>1</sub> | 1.36 ± 0.01 | (1.35, 1.37) | 1.4 ± 0.2 | (1.2, 1.6) | 0.51 ± 0.03 | (0.48, 0.54) |
| I<sub>2</sub> to U | ns | ns | 0.06 ± 0.01 | (0.05, 0.07) | 0.075 | (0.07, 0.08) |
| I<sub>2</sub> to I<sub>1</sub> | 0.015 ± 0.001 | (0.014, 0.016) | ns | ns | ns | ns |
| **While bound** |  |  |  |  |  |  |
| NG4 to I<sub>1</sub>G4 | 23.3 | (10.8, 68.8) | 22.7 | (8.25, 961) | 7.11 | (2.33, 71.1) |
| I<sub>1</sub>G4 to UG4 | 0.47 | (0.46, 0.48) | 0.41 | (0.35, 0.47) | 2.2 ± 0.5 | (1.7, 2.7) |
| **Binding** |  |  |  |  |  |  |
| U+G4 | 29 | (29, 29.8) | 32.2 | (12.9, 1460) | 30.3 | (19.4, 145) |
| N+G4 | 0.035 | (0.03, 0.04) | 0.02 | (0.01, 0.02) | 0.02 | (0.01, 0.02) |
| **Dissociation** |  |  |  |  |  |  |
| UG4 | ns | ns | ns | ns | ns | ns |
| NG4 | ns | ns | ns | ns | ns | ns |

\* All units for rates are in s<sup>-1</sup>. Binding unit is in μM<sup>-1</sup> s<sup>-1</sup>. After using 0.95 as $\chi^2$ threshold limit, error of all well-constrained rates are reported using 0.99 as $\chi^2$ threshold at boundary to give lower and upper limits, shown in parentheses. Cases with no listed error indicate either the best rates from fit or only lower limit can be detected from the confidence contour showing in Fig. S5, S8, and the reported range is predicted by Kintek Explorer.

